# 3D reconstructions of brain from MRI scans using neural radiance fields

**DOI:** 10.1101/2023.04.24.538160

**Authors:** Khadija Iddrisu, Sylwia Malec, Alessandro Crimi

## Abstract

The advent of 3D Magnetic Resonance Imaging (MRI) has revolutionized medical imaging and diagnostic capabilities, allowing for more precise diagnosis, treatment planning, and improved patient outcomes. 3D MRI imaging enables the creation of detailed 3D reconstructions of anatomical structures that can be used for visualization, analysis, and surgical planning. However, these reconstructions often require many scan acquisitions, demanding a long session to use the machine and requiring the patient to remain still, with consequent possible motion artifacts. The development of neural radiance fields (NeRF) technology has shown promising results in generating highly accurate 3D reconstructions of MRI images with less user input. Our approach is based on neural radiance fields to reconstruct 3D projections from 2D slices of MRI scans. We do this by using 3D convolutional neural networks to address challenges posed by variable slice thickness; incorporating multiple MRI modalities to ensure robustness and extracting the shape and volumetric depth of both surface and internal anatomical structures with slice interpolation. This approach provides more comprehensive and robust 3D reconstructions of both surface and internal anatomical structures and has significant potential for clinical applications, allowing medical professionals to better visualize and analyze anatomical structures with less available data, potentially reducing times and motion-related issues.

## 1 INTRODUCTION

Magnetic resonance imaging (MRI) is a powerful medical imaging technology that allows non-invasive examination of the internal structures and functions of the body. In recent years, 3D projections of MRI scans have become an increasingly popular tool for medical diagnosis and treatment planning, as they offer a detailed and accurate representation of the complex internal anatomy of the body [7]. However, long MRI acquisition sequences are particularly subject to motion-related artifacts due to breathing and cardiac pulsation [2]. This can lead to inaccuracies in 3D reconstruction, which can have significant consequences in a clinical setting where precise measurements are important for treatment planning and monitoring. This paper will explore the challenges of creating 3D projections from MRI scans and discuss the various methods and techniques used to address these challenges. Several theories have been proposed for 3D image reconstructions of MRI and computed tomography (CT) scans, some focusing on data mining and machine learning [4] while others focus on deep learning approaches [13].

Neural Radiance Fields (NeRF) is a recently developed method for synthesizing novel views of a scene from a set of 2D images taken from different viewpoints. This method uses a neural network to represent the radiance (ie, the color and intensity of light) at each point in a 3D space within a scene, allowing the generation of photorealistic 3D visualizations [9]. The main idea behind NeRF is to learn a continuous 5D function that maps the spatial coordinates and viewing direction of a scene point to its corresponding radiance value. This function is learned from a set of training images that are captured from different viewpoints, allowing the network to infer the radiance at each point from different perspectives. NeRF has shown promising results in generating photorealistic 3D visualizations from real-world data and has demonstrated improved performance over traditional methods such as multi-view stereo and voxel-based approaches. Despite its promising results, NeRF has some limitations, including the requirement for a dense set of input views covering the entire scene from many viewpoints, which can be challenging to acquire in the medical domain, where privacy regulations can hinder data collection. In addition, in the context of MRI scans, NeRF faces further challenges due to the high dimensionality of MRI data, low contrast or missing information in the images, and computational costs in processing these data. Several variations have been made to adopt NeRF to several applications while trying to solve its limitations. D-NeRF [10] extends NeRF to a dynamic domain, which aids in the reconstruction and rendering of novel images of objects in motion compared to static scenes in NeRF. [12] proposed modifications on NeRF to eliminate the need for known camera parameters, such as poses and intrinsics, while PixelNeRF [15] presents a learning framework architecture to train NeRF via a fully convolutional network on just a few input images to predict a continuous neural scene representation. There is growing interest in the use of NeRF based technologies in medical imaging, more specifically surgery scene monitoring [8] and endoscopy [6]. MedNeRF [1] is an adaptation of the generative radiance fields (GRAF) model [11] for rendering CT projections given a few or even a single-view X-ray in the medical domain. The approach not only synthesizes realistic images but also captures the data manifold and provides a continuous representation of how the attenuation and volumetric depth of anatomical structures vary with the viewpoint without 3D supervision. A new discriminator architecture provides a stronger and more comprehensive signal to GRAF when dealing with CT scans. This framework relies on an innovative architecture that builds from neural radiance fields and learns a continuous representation of CT scans by decoupling the volumetric depth and shape of the surface and intrinsic structures of the body from 2D CT scans.

In this paper, we introduce a novel method to reconstruct 3D projections inspired by MedNeRF due to its ability to model complex scenes with high fidelity, which we could call extended-MedNeRF. Although 3D reconstruction with NeRF seems promising, MRI scans present several challenges due to the multiple 2D slices and variable slice thicknesses typical of MRI data. To address these challenges, we will incorporate slice-level information by modifying the neural network architecture to take into account the unique characteristics of MRI slices. This will involve using a 3D convolutional neural network (CNN) to extract features from each MRI slice or modifying the graph neural network used in MedNeRF to incorporate information from adjacent slices. Secondly, we will address variable slice thicknesses by using slice interpolation to resample the slices to a common thickness before building the 3D model. Additionally, we will incorporate information from multiple MRI modalities into the MedNeRF framework to improve the accuracy and robustness of the resulting 3D models. Finally, we will train the model on the BraTS Challenge dataset which is a large dataset of MRI scans to ensure it can be generalized to a wide range of applications. However, reconstructions are meant to be used for visualization purposes or for healthy control subjects, our extended-MedNeRF cannot be meant to be a prediction of all clinical cases such as glioma, meningioma, or other types of lesions.

## 2 METHODS

The proposed approach comprises several parts, which are described in the following sub-sections as well as depicted in Figure 1:

1. Preprocessing
2. Feature extraction from input slices using a CNN architecture to provide a better input to extended-MedNeRF
3. Use of our extended-MedNeRF to generate 3D reconstruction

**Fig. 1.**
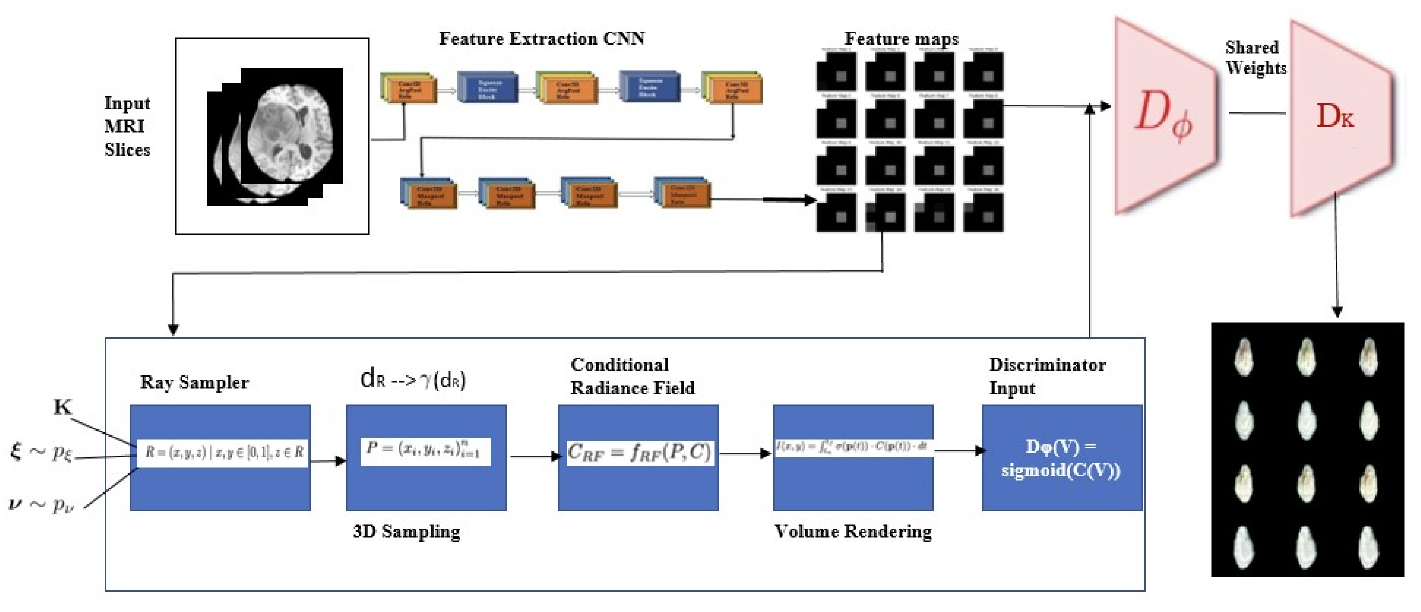
The overall architecture after preprocessing. The steps include the proposed 3D CNN, generating a feature map that highlights regions in the input image that are most important for the classification decision, and the subsequent generative radiance fields.

### 2.1 MRI Preprocessing

Generating quality results of our method on MRI slices does not come without challenges. Some of these challenges associated with the use of MRI data include low contrast and poor resolution, and we present preprocessing techniques to improve data quality. Specifically, we used image registration to correct patient movement during scans and re-scale images to improve resolution. We also propose a conditional radiance field approach, which conditions the NeRF model on additional latent codes to disentangle shape and appearance information, allowing for separate manipulation of these attributes during inference. Our experimental results demonstrate the effectiveness of our proposed approach in producing high-quality 3D reconstructions of MRI data, highlighting the potential for clinical applications in medical imaging. **Image Registration**: Aligning multiple MRI scans of the same patient can improve the quality of the data by reducing motion artifacts and allowing for better visualization of small details. Image registration improved the performance of the model by correcting for any spatial misalignments or inconsistencies in the input MRI images. By aligning the images to a common reference frame, registration can reduce artifacts and distortions in the reconstructed 3D volume, resulting in a more accurate and reliable output. Additionally, registration can also help reduce noise and improve the signal-to-noise ratio in the reconstructed volume.

#### Image Rescaling

We employed image rescaling as a preprocessing step to enhance the performance of our model. We observed that resizing the input MRI slices to a higher resolution using bicubic interpolation resulted in a significant improvement in the accuracy of our 3D reconstruction model. Bicubic interpolation is a widely used method for image rescaling [3]. The bicubic function is a two-dimensional extension of the cubic Hermite spline. Given an input image I of size *m ∗ n*, the output image *I* ’
s of size *p ∗ q* can be computed using the bicubic interpolation formula:

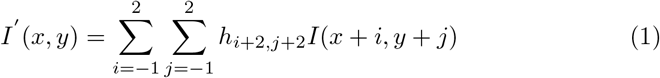

*h*_*i*+2,*j*+2_ are the bicubic coefficients, computed from the four nearest neighbour pixels to the interpolated location (x,y). The bicubic interpolation method produces smoother and more accurate results than the bilinear interpolation method, but requires more computation. This is because higher resolution images provide more detailed information about the underlying anatomy and structures, leading to more precise reconstruction of the 3D volume.

#### Contrast Enhancement

Techniques such as contrast enhancement were applied to the input images before feeding them into the network. This was done using a histogram equalization technique [14], which extends the histogram of the image to cover the entire dynamic range of the pixel values. Let *p*_*r*_(*r*_*k*_) denote the probability mass function of the input image intensities, where *r*_*k*_ represents the *k*th intensity level. The corresponding cumulative distribution function (CDF) is given by 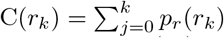.

The histogram equalization transformation function *T* (*r*_*k*_) is defined as:

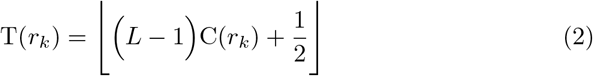

This results in a more evenly distributed pixel intensity across the image, which can help reveal subtle details that could have been hidden in the original image. By applying contrast enhancement to the input images, we were able to improve the visual quality of the reconstructed 3D projections and increase the accuracy of our model in identifying anatomical structures in the images. This is particularly important in medical imaging applications where accurate diagnosis and treatment planning depend on the ability to accurately identify and localize subtle changes in images.

### 2.2 Feature Extraction with 3D CNN

MRI data typically consists of multiple images, each representing a different slice of the 3D volume. Depending on the specific format of the MRI data, we need to do a lot of pre-processing. To adapt the baseline model on MRI slices, we used a 3D CNN for feature extraction. The 3D CNN can extract features from the MRI slices and output a 3D feature representation for each slice. To use this 3D feature representation as input to the NeRF model, we generated different views of the MRI slices from different angles. We then pass the MRI slices through the 3D CNN to extract the 3D feature representation. Next is to flatten the 3D feature representation for each MRI slice into a 1D vector, concatenate the flattened feature vectors which will result in a matrix where each row corresponds to a different slice and each column corresponds to a different feature. Our architecture consists of a 3D convolutional neural network that extracts features from the MRI slices followed by a 2D convolutional neural network to the output of the 3D CNN to generate a 2D feature representation for each MRI slice. The model comprises multiple blocks that combine a convolutional layer with an average pooling layer to reduce the size of the input feature maps while maintaining some of the original resolutions. We also perform feature recalibration by the addition of a Squeeze-and-Excitation (SE) block. The SE block is a lightweight and effective module that learns to recalibrate feature maps by adaptively weighting them based on their channel-wise importance. It consists of two operations: a squeeze operation that aggregates the spatial dimensions of each feature map into a global descriptor and an excitation operation that generates a set of channel-wise weighting coefficients. These coefficients are then multiplied with the original feature maps to obtain the recalibrated features. The features tensor generated from the CNN architecture is a 2D feature representation for each MRI slice, and can be visualized as a stack of 2D images. Each 2D image corresponds to one MRI slice and contains the learned features for that slice. We then feed the concatenated feature matrix as input to our model. The goal is to enable the model to learn to generate a 3D reconstruction of the MRI volume based on the concatenated feature matrix and the different views of the MRI slices.

### 2.3 Slice Interpolation

Slice interpolation is a commonly used technique in medical imaging to improve the resolution and quality of MRI images. It involves estimating the value of a pixel at a higher resolution based on the values of adjacent pixels using mathematical algorithms. This technique is particularly useful when the MRI slices have low resolution or when additional slices are needed to better visualize the structure or pathology of interest [5]. In our study, we employed slice interpolation not only as a preprocessing step but also as a means of generating intermediate slices between two adjacent slices in an MRI volume. This was necessary because the method we used to generate slices was completely random and we wanted to ensure that all parts of the volume were adequately sampled. We first preprocessed the MRI data by rescaling it to a common resolution, converting it to a standard file format, and applying any necessary corrections or adjustments. We then trained our model on the preprocessed MRI data to generate a 3D volume of radiance fields. However, to improve the quality of the generated images, we applied slice interpolation to generate additional slices between existing MRI slices.There are several interpolation algorithms and techniques that can be used, including linear interpolation, cubic interpolation, and Fourier interpolation. To balance the trade-off between computational resources, time, and image quality, we chose to use the cubic interpolation method, which allows us to estimate the value of a pixel at a higher resolution using a cubic polynomial based on the values of adjacent pixels. Given a set of n data points (*x*0, *y*0), (*x*1, *y*1), *·*, (*xn −* 1, *yn −* 1), the goal of cubic interpolation is to find a function f(x) that passes through each data point and approximates the underlying curve. The cubic interpolation function takes the form:

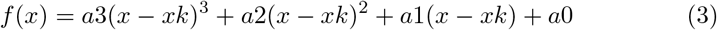

xk is the x-coordinate of the k-th data point, ak is the coefficient of the k-th term in the cubic polynomial To find the coefficients a0, a1, a2, and a3, we need to solve a system of equations that ensures that the interpolation function passes through each data point and has a continuous first and second derivative. The resulting system of equations takes the form of a tridiagonal matrix, which can be solved efficiently using standard numerical methods. Cubic interpolation is a popular choice for slice interpolation because it provides a good balance between accuracy and computational efficiency. By estimating the value of a pixel at a higher resolution using a cubic polynomial based on the values of adjacent pixels, we can generate high-quality MRI images that better represent the underlying structure and pathology of interest. In summary, slice interpolation is a useful technique for improving the resolution and quality of MRI images, and the choice of interpolation method should be carefully considered based on the specific application and data. Using slice interpolation, we were able to generate high-quality MRI images that better represent the underlying structure and pathology of interest.

### 2.4 3D Reconstruction Via extended-MedNeRF

**NeRF** is a machine learning technique that can be used for the 3D reconstruction of MRI scans. As depicted in Figure 2, NeRF uses a neural network to learn a function that maps a 3D point in space to a color and opacity, which can be used to reconstruct the 3D geometry and appearance of organs or tissues in the body from a collection of 2D MRI images. The neural network is trained on a set of images that capture the same scene from different viewpoints, and can then be used to render new views of the scene from any viewpoint. Compared to traditional methods, NeRF has the potential to produce more accurate and detailed 3D reconstructions, particularly when dealing with complex shapes or structures.

**Fig. 2.**
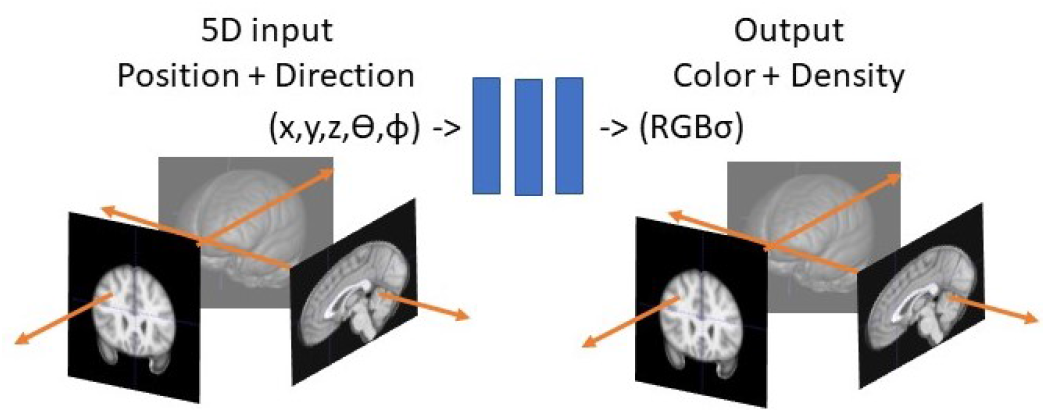
An overview of NeRF Architecture with 2D Slices of MRI scans

Adapting NeRF to MRI scans presents challenges due to differences in imaging systems, and overlapping anatomical structures in MRI data make it difficult to define edges. To address these challenges, we propose a method that adopts a generative radiance field approach. This approach employs GRAF as a backbone architecture to generate reconstructions from slices of MRI data that have undergone slice interpolation and feature extraction via NeRF.

**GRAF** is a generative model that uses a continuous representation of 3D space to generate high-quality images of scenes from any viewpoint. One of the key contributions of GRAF is its hierarchical sampling approach, which enables it to generate high-resolution images efficiently. The model also incorporates a novel attention mechanism that allows it to focus on relevant parts of the scene when generating images. The potential application of GRAF to MRI is exciting, as it could enable the generation of high-quality medical images from any viewpoint. The GRAF architecture is made up of a generator *Gθ* and a discriminator *DΦ*. Given a set of images, *Gθ* predicts a patch *P*_*pred*_, while *DΦ* compares the predicted patch with a ground-truth patch *P*_*real*_ extracted from a real image. The *K ∗ K* patch P(**u**, *s*) constituting the ray sampler of the Generator is denoted by 2D coordinates which depict the locations of pixels of the patch in the domain *Ω*

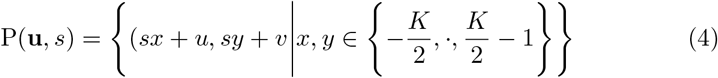

The 3D rays corresponding to the patch are determined by the patch’s image coordinates, the camera pose *ε*, and intrinsics **K**. The pixel/ray index is denoted by “r”, the normalized 3D rays by “dr”, and the number of rays by “R”. The 3D rays corresponding to the patch are determined by the patch’s image coordinates, the camera pose, and intrinsics. The pixel/ray index is denoted by “r”, the normalized 3D rays by “dr”, and the number of rays by “R”. During training, the number of rays is R = K2, where K is a constant. During inference (the process of generating the final image), the number of rays is R = WH, where W and H are the width and height of the final image. A 3D point sampling technique with stratified sampling is then applied to numerically integrate the radiance fields. Patches are then passed into a conditional radiance field, where a deep neural network maps the 3D location and viewing direction to RGB color value and volume density, conditioned on latent codes for shape and appearance. Finally, the color and density of points along a ray are used to obtain the color of the corresponding pixel, and the results of all rays are combined to generate a predicted patch in the volume renderer. The predicted patches are passed to the discriminator *DΦ*, where it is compared with a patch extracted from a real image using bilinear interpolation, allowing continuous displacements and scales while retaining high-frequency details.

Our goal is similar to [1], but in contrast, we disentangle directly in 3D MRI instead of DRR-generated CT scans. To adapt GRAF to our task of using MRI data instead of CT scans, we needed to modify the method to obtain the radiation attenuation response from MRI data. This involves sampling MRI scans to extract information on the 3D shape and appearance of anatomical structures. The density and pixel values are then computed at each sampled point using a multi-layer perceptron. The final pixel response is then obtained using a compositing operation, and the generator can predict an image patch based on the sampled MRI scans. Due to the small size of the medical datasets, the GRAF discriminator *D*(*ω*) cannot provide enough refined features to the generator *Gθ*, resulting in inaccurate volumetric estimation. Furthermore, limited training data may lead to the generator or discriminator falling into ill-posed settings, resulting in a suboptimal data distribution estimation. Classic data augmentation techniques may also mislead the generator to learn infrequent or non-existent augmented data distributions.

### 2.5 Dataset

The study employs the publicly available International Brain Tumor Segmentation (BraTS) Challenge 2021 dataset, which encompasses multi-institutional routine clinically acquired multi-parametric MRI (mpMRI) scans of glioma. The dataset comprises scans with pathologically confirmed diagnoses, along with the available MGMT promoter methylation status for glioblastoma cases with these pertinent data. Notably, the dataset has undergone augmentation since BraTS’20, with the inclusion of numerous additional routine clinically acquired mpMRI scans. To train our models, we utilized 40 slices for each of 3 patients in the 2021 dataset of the Challenge. The choice of slices was made erratically. This method was utilised to avoid bias or preconceptions about what should be included or excluded. Additionally, we performed digitally reconstructed radiograph (DRR) generation, which facilitates the removal of patient data and enables control in capture ranges and resolutions. Our training dataset consisted of 30 patients, while the remaining subjects served as the testing set.

## 3 RESULTS

We make comparisons between our model’s outcomes, two other benchmarks, and conduct a study on the impact of specific variables. Both qualitative and quantitative assessments are presented, and all models are trained for 100,000 iterations with a batch size of 8. To uniformly sample points on a sphere’s surface, we select projection parameters (u, v) that elevate slightly horizontally between 70-85 degrees and have umin = 0, umax = 1 for a complete 360-degree vertical rotation. During training, we only offer a fifth of the views (72-views each at five degrees) and allow the model to render the remaining ones.

### 3.1 Rendering From Single View MRI

The proposed reconstruction approach used has the ability to produce results of volumetric rendering which involves generating a complete 3D representation of a medical instance from a series of 2D MRI slices. However, with the use of a trained model, it is now possible to achieve this with just a single MRI slice. To achieve this, the model is trained using a relaxed reconstruction formulation, as described in [23], which involves fitting the generator to a single image. Then, the parameters of the generator are fine-tuned alongside the shape and appearance of latent vectors zs and za. In order to balance the trade-off between distortion and perception, a Mean Square Error (MSE) loss is added to the generation objective, as is common in GAN methods [24]. With this approach, it is now possible to generate a complete 3D volumetric rendering from a single MRI slice. In Figure 3 the qualitative results obtained by the proposed method are reported. This features reconstructed slices given a single MRI scan and contrast transformation of slices showing the model’s ability to differentiate between bones and other tissues by enhancing the contrast between them. This is achieved by increasing the brightness of pixels that correspond to denser structures such as bone, making them stand out more clearly from the surrounding tissue.

**Fig. 3.**
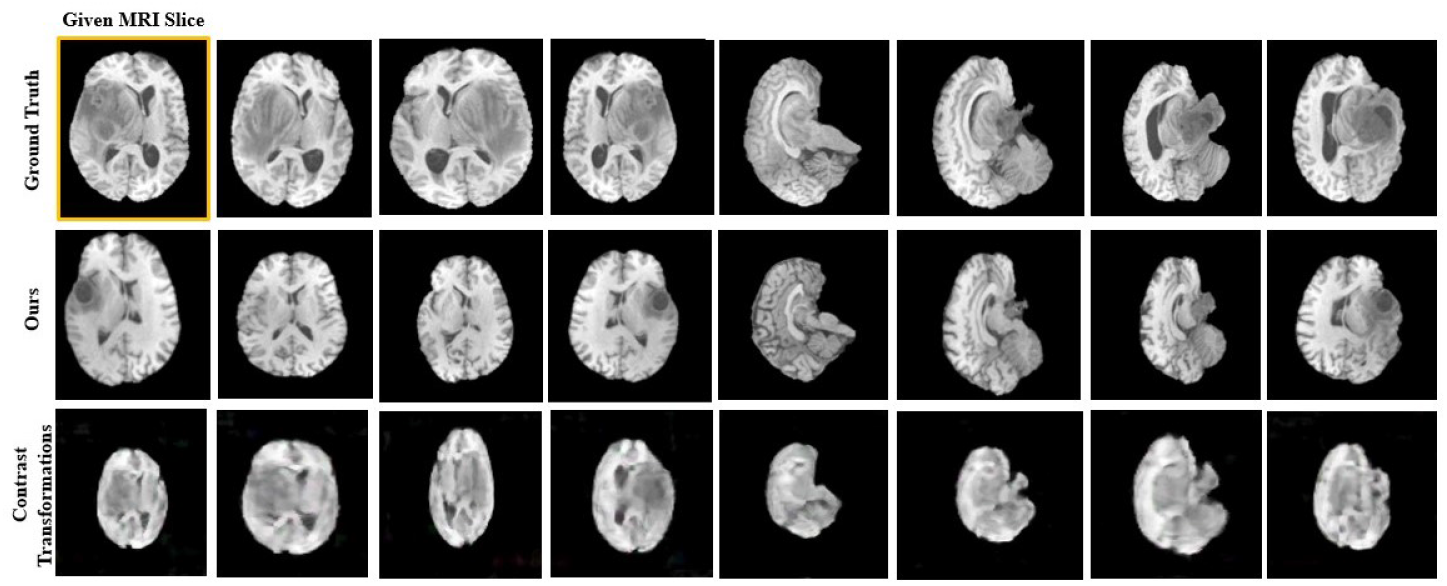
Qualitative results of MRI reconstruction

### 3.2 PSNR and SSIM

PSNR and SSIM are metrics that are used to evaluate the quality of images or videos in the field of machine learning. PSNR (Peak Signal-to-Noise Ratio) is a measure of the amount of noise present in an image or video. It compares the original image or video to a compressed or distorted version of that image or video. PSNR is defined as the ratio between the maximum possible value of a signal and the mean squared error (MSE) of the distorted signal, expressed in decibels (dB). Higher PSNR values indicate higher image or video quality, while lower PSNR values indicate lower quality. SSIM (Structural Similarity Index) is a metric that measures the similarity between two images. It takes into account both the structural information (i.e., the organization of pixels in the image) and the luminance and contrast information of the image. SSIM values range from -1 to 1, with 1 indicating perfect similarity between the two images and -1 indicating complete dissimilarity. In the proposed experiments, the PSNR and SSIM were, respectively, 25.01 *±* 1.17 and 0.879 *±* 0.07.

### 3.3 FID and KID with Other Models

FID measures the similarity between two datasets of images, one generated by a generative model and the other real, using feature statistics calculated from a pre-trained neural network. Specifically, FID measures the distance between the feature distributions of the generated images and the real images. Lower FID scores indicate higher quality generated images that are more similar to real images. KID also measures the similarity between two datasets of images, but instead of using feature statistics, it uses kernel-based methods to compare the distributions of the images. KID measures the distance between the two distributions of feature representations from the generated images and the real images. Lower KID scores also indicate higher quality generated images that are more similar to real images. More specifically, the proposed experiments result in a FID and KID score, respectively, of 160.12 and 0.16 *±* 0.003.

## 4 DISCUSSIONS

A 3D model of a scene can be created from a collection of 2D images using the computer vision method known as structure from motion (SfM). SfM is a multi-step process that includes point-cloud reconstruction, feature recognition and matching, and camera pose estimation. Recent advances in deep learning have led to a new method, called NeRF, to create photo-realistic 3D scenes from a collection of 2D images. NeRF has produced stunning outcomes when producing high-fidelity 3D models in several fields, and its adoption by the medical imaging community is also growing. In this paper we explored the idea of reconstructing an MRI volumes from heterogeneous slices, with the ultimate goal of reducing the number of scans during an MRI session or motion artefacts related to long acquisitions at least for educational purposes. Given a data driven generation, it should be used carefully in clinical context as the reconstructed data might hide relevant specific elements in certain pathological cases, and this represents a limitation shared by many generative approaches. Nevertheless, this can lead new ways for data visualization. We called the proposed approach extended-MedNeRF. In particular, we introduced a radiation attenuation response from MRI in the generative radiance field. Moreover, we used challenging scenarios as low-sample size for training and non-uniform slicing. The reconstruction produced promising results that, even if they cannot be used for clinical evaluation, opens a new way to reduce MRI acquisitions and consequent motion corrections. Future work includes tests on brain scans from people of different ages and other MRI modalities.

## Acknowledgments

This publication is partially supported by the European Union’s Horizon 2020 research and innovation programme under grant agreement Sano No 857533, and partially by the International Research Agendas programme of the Foundation for Polish Science, co-financed by the European Union under the European Regional Development Fund.

## Footnotes

1 http://braintumorsegmentation.org

